# Mating Strategies of Invasive Versus Indigenous Crayfish: Multiple Paternity as Driver for Invasion Success?

**DOI:** 10.1101/2021.05.28.445155

**Authors:** Caterina Francesconi, Mălina Pîrvu, Anne Schrimpf, Ralf Schulz, Lucian Pârvulescu, Kathrin Theissinger

**Author notes:** Authors having equal contribution (Lucian Pârvulescu, Kathrin Theissinger). Corresponding author: Kathrin Theissinger.

## Abstract

The invasive spiny-cheek crayfish (*Faxonius limosus*) has been able to colonize many European waterbodies since its first introduction into Europe, threatening the indigenous crayfish fauna. *Faxonius limosus*’ remarkable reproductive plasticity has been suggested as an important factor contributing to this species’ alarming invasiveness. This is the first study comparing the reproductive strategies of an invasive (*F. limosus*) and a sympatric indigenous crayfish (*Pontastacus leptodactylus*). We investigated if and how parthenogenesis and multiple paternity contribute to the invasion process in the River Danube. Using microsatellites, we genotyped the offspring and their mothers of 11 clutches of *F. limosus* and 18 clutches of *P. leptodactylus*. While no parthenogenesis has been found in *F. limosus*’ populations, multiple paternity has been detected for the first time in both species, with comparable incidence. The results of the study indicate that multiple paternity does not play a dominant role in *F. limosus*’ successful colonization of the Danube. However, the presented results have to be regarded as pilot study, with a limited number of samples and loci investigated. Given the relevance of mating system knowledge for management measures, future studies with larger sample number could provide precious contributions to the conservation actions.

## INTRODUCTION

Since the mid-19^th^ century, at least 11 non-indigenous crayfish species (NICS) have been purposefully or accidently introduced into Europe (Jussila *et al*. 2015) and became invasive (Holdich *et al*. 2009); (Jussila *et al*. 2015). Today NICS are widespread across Western and Central Europe and they keep spreading (Kouba *et al*. 2014), causing changes in the distribution patterns of the indigenous crayfish’s populations (Jussila *et al*. 2015). Crayfish represent keystone species in freshwater ecosystems, influencing the food web by being an important resource for other animals and by feeding on local vegetation and invertebrates (Reynolds *et al*. 2013). Several NICS can yield high-density populations and, even when replacing indigenous crayfish, they can have a significant effect on the indigenous biota such as benthic fish, molluscs, and macrophytes (Gherardi, 2007). Therefore, the consequences of their introduction, ranging from the danger posed to indigenous crayfish species to the stress imposed on the ecosystem structure, cannot be overlooked (Hobbs *et al*. 1989; Momot, 1995; Reynolds *et al*. 2013).

The North American spiny-cheek crayfish *Faxonius limosus* (Rafinesque, 1817) was the first non-indigenous crayfish species introduced into Europe in 1890 (Holdich *et al*. 2009; Filipová *et al*. 2011). Since its first introduction into Europe this invasive species has been able to quickly increase its population and successfully colonize many European waterbodies and can now be found in at least 25 European countries (Kouba *et al*. 2014; Trichkova *et al*. 2015; Govedič, 2017; Kaldre *et al*. 2020). Many studies have addressed the alarming invasiveness of *F. limosus* (e.g. Kozák *et al*. 2006; Buřič *et al*. 2013; Kouba *et al*. 2014; Pârvulescu *et al*. 2015). The spread of this invasive species represents a threat to all indigenous crayfish (Holdich and Pöckl, 2007). This includes direct stress by competing for the same resources (Holdich and Pöckl, 2007; Lele and Pârvulescu, 2017; Pacioglu *et al*. 2020) and by a superior aggressive behaviour (Lele and Pârvulescu, 2017; Weis, 2010; Pârvulescu *et al*. 2021), and indirect stress by acting as vector for the crayfish plague pathogen *Aphanomyces astaci* (Holdich and Pöckl, 2007).

*Faxonius limosus* exhibits r-selected life strategies typical for invasive crayfish species, such as short life cycle, high fecundity, early maturation, and capability of taking maximum advantage from abundant resources (Lindqvist and Huner, 1999; Kozák *et al*. 2007). This species possesses an exceptional reproductive plasticity (Buřič *et al*. 2013). The division of the mating period in spring and autumn seasons maximizes the probability of successful mating (Buřič *et al*. 2013). The ability of both females and males to alternate between sexually active and inactive forms allows them to direct the resources utilization toward structures useful for specific life stages (Buřič *et al*. 2010b; Buřič *et al*. 2010a). *Faxonius limosus* is also capable of increasing its fecundity, leading to a quick growth of the population by maximizing the exploitation of the resources made available by the decrease of indigenous crayfish populations (Pârvulescu *et al*. 2015). *Faxonius limosus* has been found capable of facultative parthenogenesis under laboratory conditions (Buřič *et al*. 2011; Buřič *et al*. 2013). The occurrence of parthenogenesis in the wild would allow reproduction under suboptimal conditions, facilitating the colonization of new habitats from small founding populations (Buřič *et al*. 2011). Lastly, long-term sperm storage (Buřič *et al*. 2013) also allows the reception of spermatophores from different males, while enabling this animal to circumvent adverse environmental conditions (Walker *et al*. 2002). The potentially resulting multiple paternity can enhance the success of a population introduced in a new environment by increasing its genetic diversity.

Multiple paternity is widely spread both among vertebrates and invertebrates (Jennions and Petrie, 2000; Avise *et al*. 2011), with variable incidence across and within species (Taylor *et al*. 2014). Multiple mating is associated with considerable costs for females, such as loss of time and energy, and increased risk of predation, diseases and injuries (Slatyer *et al*. 2012). Nonetheless, the ubiquitous nature of this reproductive strategy indicates that multiple paternity provides significant benefits (Hosken and Stockley, 2003). One of the major genetic benefits is the increased heterozygosity (Holman and Kokko, 2013; Taylor *et al*. 2014). High heterozygosity is linked to disease resistance, greater development stability, competitiveness, hatchability and survivorship, and the ability to respond to novel biotic and abiotic stimuli (Holman and Kokko, 2013; Taylor *et al*. 2014). Therefore, multiple paternity is an efficient method to increase offspring fitness (Palmer and Oldroyd, 2000; Taylor *et al*. 2014) and effective population size (Zeng *et al*. 2017). Multiple paternity has also been linked to the invasion success of non-indigenous species in reptiles (Eales *et al*. 2010), fishes (Zeng *et al*. 2017), mammals (Miller *et al*. 2010), gastropods (Le Cam *et al*. 2009; Rafajlović *et al*. 2013), insects (Ding *et al*. 2017) and malacostracans (Yue *et al*. 2010).

Many studies addressing the invasiveness of crustaceans compare indigenous and invasive species’ behaviours (Weis, 2010). This approach allows for a deeper understanding of the contingency of a successful invasion (Stohlgren and Schnase, 2006; Weis, 2010). However, there is a general lack of comparative studies regarding the reproductive strategies of indigenous *vs*. invasive species (Weis, 2010). In this pilot study we analysed and compared, for the first time, the reproductive strategies of an invasive (*F. limosus*) and an indigenous (*Pontastacus leptodactylus* Eschscholtz 1823) crayfish species occurring in sympatry in the river Danube. *Faxonius limosus* has been detected in the Romanian Danube for the first time in 2009 after downstream dispersal from Serbia (Pârvulescu *et al*. 2009) and has partially replaced the indigenous *P. leptodactylus* in the upper Romanian Danube (Pârvulescu *et al*. 2012; Pârvulescu *et al*. 2015). We aimed to understand the role of different reproductive strategies in the success of the invasion by genotyping female crayfish and their offspring using microsatellite markers. Firstly, we investigated the presence of parthenogenesis in wild populations of *F. limosus*. Secondly, we analysed the incidence of multiple paternity in both crayfish species. We hypothesized that parthenogenesis and multiple paternity are prominent in the invasive species, thus acting as additional drivers for its invasion success.

## MATERIALS AND METHODS

### Sampling and study design

Female crayfish with eggs were collected in the upper Danube during summer 2016 (Fig. 1). The river was divided into three sectors, based on the time-line of the invasion and the coexistence state of the two species: an old-invaded sector (OID) where the invasion of *F. limosus* dates back at least ten years; an active invasion front sector (IF) where both species coexist and *F. limosus* has been present for no more than three years at the time of samples collection; a non-invaded sector (NID) where only *P. leptodactylus* is present (Fig. 1). The crayfish were caught in the littoral area using bait-trap left overnight. In total, eleven female *F. limosus* and their clutches were collected for the analysis (Tab. 1). Four of them were sampled from the IF sector and seven from the OID sector. Six female *P. leptodactylus* and their clutches were sampled from each sector of the Danube river. Three families consisted of mother and embryos extracted from eggs, while in the other clutches (N = 15) the mothers were collected with juveniles (Tab. 1). Tissue samples were stored in 96% ethanol. The sampling was conducted according to animal welfare regulations, permits were obtained from the Iron Gates Nature Park Administration and the Regional Environmental Protection Agency.

**Fig. 1.**
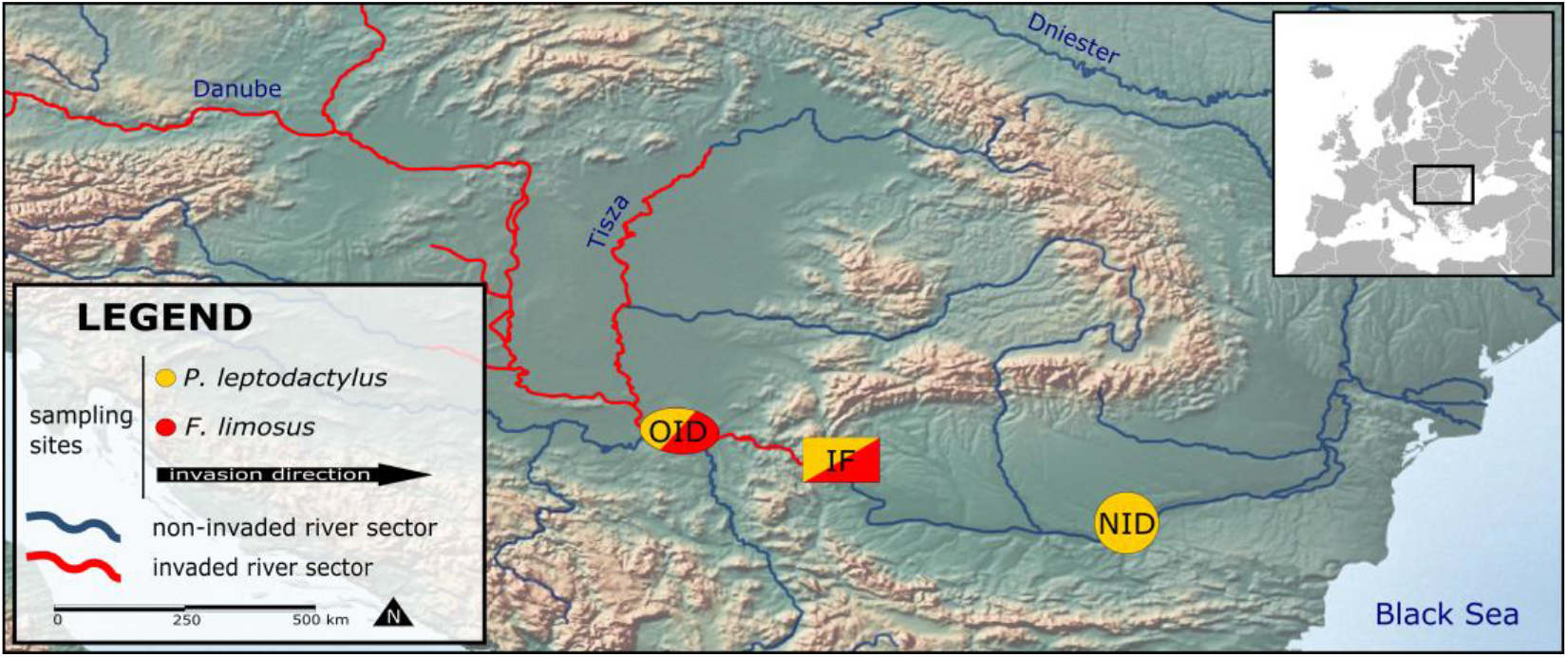
Sampling sites along the river Danube. NID, non-invaded sector; OID, old-invaded sector; IF, invasion front.

**Tab. 1.** Sampling location, sample size, offspring stage and inferred genetic paternity in 18 clutches of *Pontastacus leptodactylus* and 11 clutches of *Faxonius limosus*.

| Species | Sector | Mother's ID | Known maternal genotype | Offspring stage | No. of offspring | No. of alleles | No. of fathers | MP |
| --- | --- | --- | --- | --- | --- | --- | --- | --- |
| PL | NID | E06 | Yes | Fertilized eggs | 17 | 8 | 1 | No |
| PL | NID | E22 | No | Fertilized eggs | 20 | 7 | 1 | No |
| PL | NID | F05 | No | Juveniles | 18 | 7 | 1 | No |
| PL | NID | F11 | Yes | Juveniles | 17 | 7 | 1 | No |
| PL | NID | F28 | No | Juveniles | 18 | 8 | 1 | No |
| PL | NID | F29 | No | Juveniles | 19 | 7 | 2 | Yes |
| PL | OID | SRB061 | Yes | Juveniles | 18 | 8 | 1 | No |
| PL | OID | SRB111 | Yes | Juveniles | 15 | 8 | 2 | Yes |
| PL | OID | SRB141 | Yes | Juveniles | 19 | 8 | 1 | No |
| PL | OID | SRB171 | Yes | Juveniles | 19 | 8 | 1 | No |
| PL | OID | SRB181 | Yes | Juveniles | 18 | 8 | 1 | No |
| PL | OID | SRB204 | Yes | Juveniles | 7 | 4 | 1 | No |
| PL | IF | ESL1 | Yes | Juveniles | 10 | 8 | 1 | No |
| PL | IF | ORS011 | Yes | Juveniles | 17 | 8 | 1 | No |
| PL | IF | ORS021 | Yes | Juveniles | 14 | 8 | 1 | No |
| PL | IF | ORS031 | Yes | Juveniles | 8 | 8 | 1 | No |
| PL | IF | ORS041 | Yes | Juveniles | 6 | 8 | 1 | No |
| PL | IF | ORS254 | Yes | Fertilized eggs | 22 | 8 | 1 | No |
| FL | IF | FL0036 | Yes | Juveniles | 11 | 6 | 3 | Yes |
| FL | IF | FL0049 | No | Juveniles | 8 | 6 | 1 | No |
| FL | IF | FL0072 | No | Juveniles | 7 | 6 | 1 | No |
| FL | IF | FL0268 | Yes | Juveniles | 9 | 6 | 1 | No |
| FL | OID | FL0001 | Yes | Juveniles | 11 | 6 | 3 | Yes |
| FL | OID | FL0085 | Yes | Juveniles | 9 | 6 | 1 | No |
| FL | OID | FL0098 | No | Juveniles | 4 | 6 | 1 | No |
| FL | OID | FL0111 | No | Juveniles | 8 | 6 | 1 | No |
| FL | OID | FL0124 | Yes | Juveniles | 4 | 6 | 1 | No |
| FL | OID | FL0137 | No | Juveniles | 6 | 6 | 1 | No |
| FL | OID | FL0155 | Yes | Juveniles | 9 | 6 | 1 | No |

MP, multiple paternity; PL, *Pontastacus leptodactylus*; FL, *Faxonius limosus*; NID, non-invaded sector; OID, old-invaded sector; IF, invasion front.

### DNA isolation and microsatellites genotyping

Maternal DNA from female pereopod tissues and progeny DNA from whole eggs or half juveniles was extracted using a modified high salt DNA extraction protocol (Aljanabi and Martinez, 1997). DNA pellets were eluted in 50 μL 1xTE Buffer.

For *P. leptodactylus* the genotyping was carried out using eight microsatellites loci described in Gross *et al*. (2017). The Type-it^®^ Microsatellites PCR kit (QIAGEN) was used to amplify all loci across two multiplex PCRs: multiplex A consisted of Aast4_5, Aast4_12, Aast4_32 and multiplex B consisted of Aast4_8, Aast4_16, Aast4_26, Aast4_30, Aast4_34. The PCR reaction (6 μL) contained: 0.5 μL of Q-Solution, 2.5 μL of Type-it Multiplex PCR Master Mix, 2 μL of Primer mix (0.25 μL of each primer), 1 μL of RNase-free water and 1 μL of template DNA. The PCR protocol was the following: initial denaturation step (5 min, 95°C); 30 cycles of denaturation (30 sec, 94°C), annealing (90 sec, 57°C) and extension (60 sec, 72°C); final extension step (30 min, 60°C).

To genotype *F. limosus* six microsatellite primers were chosen following Buřič *et al*. (2011). The primers were grouped in three PCRs. Multiplex A included PclG_08, 2.12 and 3.1, multiplex B included PclG_02, PclG_26 and a single PCR was run for PclG_37. The PCR reaction was conducted under the same conditions as for *P. leptodactylus* with the exception of the annealing temperature for primers PclG_08 and PclG_37, set at 53°C. The fragment lengths were detected on a Beckman Coulter CEQ™ 8000 system. The raw data was analysed using the software GeneMarker v2.6.4 (Soft-Genetics, State College, PA, USA).

### Data analyses

For *F. limosus*, genotypes of the eleven females and four additional adults sampled in the same survey were used to calculate population allele frequencies and to verify lack of linkage disequilibrium and agreement with the Hardy-Weinberg equilibrium. For *P. leptodactylus* allele frequencies were calculated using only 15 of the 18 females analysed, as it was not possible to genotype the mothers from three clutches due to missing data. For both species allele frequencies, presence of linkage disequilibrium and agreement with the Hardy-Weinberg equilibrium of the adult crayfish were tested with GENEPOP 4.2 (Raymond and Rousset, 1995). The presence of null alleles per locus was tested with Micro-Checker (van Oosterhout *et al*. 2004). A genotyping error rate of 0.05 for *F. limosus* and 0.04 for *P. leptodactylus* was calculated by blindly repeating 10% of the samples per species. The occurrence of parthenogenesis in *F. limosus* was verified by comparing the mothers’ genotypes to the genotypes of the respective offspring. Parthenogenesis was assumed if all offspring had the identical multilocus genotype. Parentage analysis for both species was conducted using the software GERUD2.0 (Jones, 2005), which reconstructs for each progeny arrays the minimum number of fathers and their genotypes using polymorphic, codominant markers. The software ranks the parental genotype combinations by likelihood. The mother’s genotype is not needed, but the offspring array has to share the same mother. GERUD2.0 does not accept missing data, therefore specimens with too much missing data were not considered for the analysis. Chi-square test was used to infer the significance of the different incidence of multiple paternity between the two crayfish species.

The software PrDM (Neff and Pitcher, 2002) was used to evaluate the probability of detecting multiple paternity (PrDM) based on the number of alleles per locus and their frequency, the number of samples and sires’ contribution. Eight different scenarios were used to represent situations of equal males’ contribution (50:50, 33.3:33.3:33.3 and 25:25:25:25), moderate skewed contribution (66.7:33.3 and 57:28.5:14.5) and highly skewed contribution (70:10:10:10, 90:10 and 85:5:5:5). Those scenarios were based on recommendations by Neff and Pitcher (2002) and on available multiple paternity data on crayfish (Walker *et al*. 2002; Yue *et al*. 2010). The average number of offspring per clutch was used for those simulations. The resulting values represent the power of the applied genetic markers. Higher values indicate higher probabilities to detect multiple paternity in the described scenario.

## RESULTS

### Pontastacus leptodactylus

No linkage disequilibrium was detected, and the population was in Hardy-Weinberg equilibrium. No evidence of null alleles was found. Locus Aast4_32 was monomorphic in our population. Excluding locus Aast4_32, the expected heterozygosity of the seven remaining loci varied from 0.14 to 0.61, with an average value of 0.49 (Tab. 2). The observed heterozygosity ranged from 0.15 to 0.67, with an average value of 0.47. Excluding locus Aast4_32, the mean number of alleles per locus was 2.57. Locus Aast4_26 was not genotyped successfully in some families (F29, F11, F05 and E22, S1 Appendix) due to missing data and was therefore excluded from the parentage analyses for the respective clutches. For the family SRB204 the parentage analysis was conducted using only four loci due to missing data from loci Aast4_12, Aast4_32, Aast4_26, Aast4_34 (S1 Appendix). For clutches E22, F05, F28 and F29 the paternity analysis was conducted with unknown mother genotype due to missing data.

**Tab. 2.** Characterization of microsatellite loci for *Faxonius limosus* and *Pontastacus leptodactylus*. Estimates are based on adult female genotypes.

| Species | Locus | N | Na | Ho | He |
| --- | --- | --- | --- | --- | --- |
| FL | 3.1 | 11 | 3 | 0.8182 | 0.6405 |
| FL | 2.12 | 15 | 1 | 0.0000 | 0.0000 |
| FL | PclG_08 | 14 | 2 | 0.7143 | 0.4592 |
| FL | PclG_02 | 14 | 1 | 0.0000 | 0.0000 |
| FL | PclG_26 | 12 | 3 | 0.7500 | 0.5799 |
| FL | PclG_37 | 11 | 7 | 0.6364 | 0.8140 |
| PL | Aast4_12 | 15 | 3 | 0.5333 | 0.5800 |
| PL | Aast4_32 | 15 | 1 | 0.0000 | 0.0000 |
| PL | Aast4_5 | 15 | 3 | 0.5333 | 0.5800 |
| PL | Aast4_16 | 15 | 4 | 0.6000 | 0.6067 |
| PL | Aast4_34 | 13 | 3 | 0.1538 | 0.1450 |
| PL | Aast4_8 | 15 | 3 | 0.6667 | 0.4867 |
| PL | Aast4_26 | 14 | 3 | 0.4286 | 0.5383 |
| PL | Aast4_3 | 15 | 2 | 0.4000 | 0.4800 |

PL, *Pontastacus leptodactylus*; FL, *Faxonius limosus*; n, sample size; Na, number of alleles; Ho, observed heterozygosity; He, expected heterozygosity.

Altogether, 284 offspring were successfully genotyped. The number of genotyped progeny per clutch ranged from six to 22 (with an average of 15.8). The PrDM simulations produced probabilities ranging from 0.438 to 0.955 (Tab. 3). The PrDM value of 0.438 was obtained with the 90:10 scenario, while all other probabilities were higher than 0.622. Multiple paternity was detected in two clutches (11%), F22 and SRB111, respectively sampled in the non-invaded sector and the old-invaded sector (Tab. 1). The minimum number of fathers for both clutches was two. The mean number of sires per brood was 1.1.

**Tab. 3.** Probability of detecting multiple paternity (PrDM) in *F. limosus* and *P. leptodactylus* using 6 and 8 microsatellite loci respectively. Three mating scenarios are tested: even contribution from the males, moderately skewed contribution, and highly skewed contribution toward one male. Two to four males were taken into consideration.

| Number of males | Mating scenario<br>(paternal skew) | Number of offspring |  |
| --- | --- | --- | --- |
|  |  | *P. leptodactylus* | *F. limosus* |
|  |  | 16 | 8 |
| 2 | 50:50 | 0.717 | 0.756 |
|  | 66.7:33.3 | 0.698 | 0.718 |
|  | 90:10 | 0.438 | 0.362 |
| 3 | 33.3:33.3:33.3 | 0.903 | 0.904 |
|  | 57:28.5:14.5 | 0.856 | 0.841 |
| 4 | 25:25:25:25 | 0.955 | 0.945 |
|  | 70:10:10:10 | 0.838 | 0.769 |
|  | 85:5:5:5 | 0.622 | 0.516 |

### Faxonius limosus

No linkage disequilibrium among microsatellite loci was detected. Two loci (2.12 and PclG_02) were monomorphic in our population. No null alleles were detected. All loci, with the exception of locus PclG_37, were in Hardy-Weinberg equilibrium. Excluding loci 2.12 and PclG_02, the expected heterozygosity varied from 0.46 to 0.81 for the four remaining loci with an average value of 0.62, while the observed heterozygosity ranged from 0.64 to 0.82 with an average value of 0.73 (Tab. 2). Excluding the two monomorphic loci, the mean number of alleles per locus was 3.75. Locus PclG_24 was excluded from all analysis due to too much missing data.

A total number of 86 offspring was successfully genotyped for all six loci. The number of analysed progeny per clutch ranged between four and 11 (with an average of 7.8). No parthenogenesis was detected in the clutches, as the offspring genotypes exhibited paternal alleles and were no clones from their mothers (S1 Appendix). For clutches FL0049, FL0072, FL0098, FL0111 and FL0137 the paternity analysis was conducted with unknown mother genotype due to missing data in some loci (S1 Appendix). The PrDM ranged from 0.362 and 0.945 (Tab. 3). The lowest probabilities were obtained in scenarios with high skewed paternal contribution (90:10 and 85:5:5:5). All the other probabilities are greater than 0.718. In total, two clutches (18%), one sampled in the OID sector and on in the IF sectors, were sired by multiple males (Tab. 1). Chi-square test indicated no significant difference in the incidence of multiple paternity in *F. limosus* and *P. leptodactylus*.

## DISCUSSION

To our knowledge, this is the first study investigating the reproductive strategies of an invasive (*F. limosus*) and an indigenous (*P. leptodactylus*) crayfish species occurring in sympatry. No parthenogenesis has been found in *F. limosus*’ populations. Nonetheless, the hypothesis of *F. limosus* resorting to this mating strategy in the wild still needs to be further investigated. The demonstrated capability of this crayfish of reproducing asexually under laboratory conditions (Buřič *et al*. 2011; Buřič *et al*. 2013) and the parthenogenesis documented in one wild population of another crayfish species (Martin *et al*. 2007) provide a sound theoretical background for this hypothesis.

In this study, multiple paternity has been documented in both *F. limosus* and *P. leptodactylus* for the first time. This mating strategy has been detected in 18% of the clutches of the invasive *F. limosus* and in 11% of the clutches of the indigenous *P. leptodactylus*, but the difference in the incidence of multiple paternity between the two species was not significant. However, our results are not conclusive, as the study has important limitations. The chosen markers show low genetic diversity for our populations, with three monomorphic loci across the two species and a low mean number of alleles per locus. Therefore, the resolution was too low to adequately address the research questions. Due to lack of project time and funding, it was not possible to use different markers after the initial analysis. The study was further limited by the small sample size for *F. limosus*, both in terms of number of clutches and number of offspring per clutch. The study was originally designed to include at least 20 offspring per clutch, but the sample size had to be reduced due to the impossibility of producing data for many loci of an elevated number of specimens. All these factors contributed to a low probability of detecting multiple paternity with our genetic markers and dataset.

Multiple paternity has been already associated with successful biological invasions and colonization of new habitats in a variety of animals, both in vertebrates (Eales *et al*. 2010; Miller *et al*. 2010; Ding *et al*. 2017) and invertebrates (Le Cam *et al*. 2009; Rafajlović *et al*. 2013; Zeng *et al*. 2017), including crayfish (Yue *et al*. 2010). It is suggested that multiple paternity leads to higher hatchability and greater early stage survivorship, probably due to avoidance of genetic incompatibility (Zeh, 1997; Tregenza and Wedell, 1998; Newcomer *et al*. 1999). This polyandrous mating strategy, where one female mates with several males, is associated with increased heterozygosity in the offspring (Holman and Kokko, 2013; Taylor *et al*. 2014). As a result, multiple paternity may act as a buffer against the loss of genetic diversity caused by the bottleneck event that may have occurred after the invasive species’ introduction. Multiple paternity has been linked to increased fertility as it could ensure an adequate sperm supply, guaranteeing the fertilization of all the eggs produced by the female (Birkhead and Pizzari, 2002; Hosken and Stockley, 2003). Such benefit could be particularly relevant, as a higher production of eggs by females of *F. limosus* has been observed in the newly invaded habitat (Pârvulescu *et al*. 2015). Females of *F. limosus* can store viable sperm for several months (Buřič *et al*. 2013). The combination of this feature with multiple mating could increase its initial effective population size in the invaded environment (Eales *et al*. 2010). All of those benefits linked to multiple paternity, when associated with *F. limosus* fast life cycle, could be decisive in the establishment of a fast growing, quickly adapting population.

Finally, the importance of acquiring deeper knowledge of *F. limosus* mating behaviours lies in the repercussion it may have both on the management of the invasive *F. limosus* and the conservation of indigenous crayfish species (e.g. Sutherland, 1998; Caro, 1999; Rogowski *et al*. 2013; Wildermuth *et al*. 2013). Management of invasive crayfish species has often produced less than encouraging results (Hyatt, 2004; Freeman *et al*. 2010; Gherardi *et al*. 2011). Part of this inefficiency (besides other reasons, like the difficulty of catching small crayfish) may be traced back to a lack of relevant knowledge on mating behaviours and life strategies of the targeted species (Rogowski *et al*. 2013). In a successful management plan for control or removal of invasive species, the rate of removal must exceed the growth of the population (Bomford and O’Brien, 1995). Therefore, the use of models to estimate population trends is essential (Kajin *et al*. 2012). The predictive value of those demographic models can be enhanced by incorporating information on the mating system (Wildermuth *et al*. 2013). This could be especially useful for species with considerable reproduction ability and plasticity such as crayfish. For these reasons, we believe that further studies should be conducted on *F. limosus*’ mating strategies and on their comparison with the mating strategies of indigenous crayfish. While this study did not lead to significant results, it highlighted critical issues met while inferring *F. limosus*’ and *P. leptodactylus*’ mating systems. Overcoming the limitations of this study by choosing markers with higher polymorphism and bigger sample sizes would lead to more reliable results that could be useful for management plans.

## SUPPLEMENTARY FILE

The dataset supporting the conclusions of this article is presented in Supplementary material.

## ACKNOWLEDGEMENTS

This work was supported by a grant of the Romanian National Authority for Scientific Research and Innovation, UEFISCDI, project number PN-II-RU-TE-2014-4-0785. We would like to thank Britta Wahl-Ermel for helping with the samples analysis and Mišel Jelić for helping with the selection of the markers.

